# Lineage-specific microbial protein prediction enables large-scale exploration of protein ecology within the human gut

**DOI:** 10.1101/2024.05.29.596415

**Authors:** Matthias Schmitz, Nicholas J. Dimonaco, Thomas Clavel, Thomas C.A. Hitch

## Abstract

Microbes use a range of genetic codes and gene structures, yet these are ignored during metagenomic analysis. This causes spurious protein predictions, preventing functional assignment which limits our understanding of ecosystems. To resolve this, we developed a lineage-specific gene prediction approach that uses the correct genetic code based on the taxonomic assignment of genetic fragments, removes partial predictions, and optimises prediction of small proteins. Applied to 9,634 metagenomes and 3,594 genomes from the human gut, this approach increased the landscape of captured expressed microbial proteins by 78.9%, including previously hidden functional groups. Optimised small protein prediction captured 3,772,658 small protein clusters, many with antimicrobial activity. Integration of the protein sequences and sample metadata into a tool, InvestiGUT, enables association of protein prevalence with host parameters. Accurate prediction of proteins is critical for understanding the functionality of microbiomes, hence this work will enhance understanding mechanistic interactions between microbes and hosts.

## Introduction

Microbiome analysis has focused on the study of taxonomic groups, although it is the functionality of these taxa that is of interest^1,2^. Microbes that perform the same function, or work together to form a functional group, are referred to as members of the same ‛guild’^3^. Guild-based analysis of metagenomic datasets provides some functional insight, but represents an indirect method of studying functionality, as the unit of study remains the taxa, rather than directly studying the function of interest. Such inference of functionality from taxonomy is reductionist given the variability between strains of the same species, thereby expanding the functional capacity of microbial communities^4^.

Methods derived from functional ecology^4–9^ can be applied to the study of proteins and their functions. We term this ‛protein ecology’, which aims to study the ecological distribution of proteins or functions as the unit of study ^6,10^. The direct study of proteins/functions takes into account horizontal gene transfer, which can lead to the sharing of the protein of interest outside the taxa being studied, although transcriptional activity and genomic context can modify the functionality of a protein^11–13^. It has been shown that horizontal gene transfer occurs at high rates in the human gut microbiome, particularly in industrialised populations^14^, facilitating the spread of antibiotic resistance genes, virulence factors and other traits that affect the progression of human disease^15^.

Computational methods such as AnnoTree have been developed to study the distribution of functions across the microbial tree of life, highlighting functional variation between taxa^16^. Studying the ecology of proteins, rather than species, can provide insights into the ecological importance of protein-bearing species and the ecological pressures that drive protein evolution^17^. In the context of the human gut microbiome, species-independent functional studies have provided insights into the association of bile acid and sulphur metabolic genes with colorectal cancer^18^, and type 3 secretion system effectors with human health conditions^19,20^.

Gene catalogues also place the proteins/genes as the unit of study by providing a reference source of non-redundant genes for either descriptive studies of a microbiome^21^, or targeted analysis of microbial functionality^22,23^. So far, each gene catalogue has ignoring the diversity of genetic codes used by bacteria^24–26^ and the existence of eukaryotic multiple exon genes^27^. This is indicative of the trend in human gut microbiota research to focus on prokaryotes, neglecting other taxa^28^. This is detrimental as eukaryotes and viruses directly influence human health, including via immune modulation^29^.

We present a workflow using the taxonomy of metagenomic contigs to inform protein sequence prediction. Application of this approach to the human gut uncovered a multitude of previously missed proteins and provided an improved gene catalogue. To facilitate protein ecology studies of the human gut, we developed InvestiGUT, a tool that identifies associations between protein sequence prevalence and host parameters.

## Methods

### Selection of optimal gene prediction tools

To test the performance of gene prediction tools for large-scale metagenomic predictions, 13 different tools (**Supplementary Table 1**) were evaluated individually as well as in combinations of two or three. Among these, three tools were designed for eukaryotic sequences ^30–32^, nine focused on prokaryotes^24–27,33–39^, and one for viruses^40^. Both Prodigal, and Pyrodigal^39^ (v2.1.0), an actively maintained successor to Prodigal^27^ (v2.6.3), were included in the comparisons. However, as Pyrodigal was determined to give identical results to Prodigal, Prodigal was removed from the comparisons. Three of the eukaryotic tools (AUGUSTUS^30^ v3.3, GlimmerHMM^31^ v3.0.4, and SNAP^32^ v2006-07-28) rely on the selection of a model before genes can be predicted. AUGUSTUS was run with the built-in saccharomyces_cerevisiae_S288C model on all eukaryotic genomes. Both other tools had no default fungal model, so a model was built based on *Saccharomyces cerevisiae* S288C (GCF_000146045.2).

The genomes and corresponding gene predictions included in the tool comparison are detailed in **Supplementary Table 2**. The bacteria studied included the six Bacteria from the original ORForise comparison, supplemented with bacterial species deemed pathogenic^41–45^ or health-associated^46–51^. In addition to the bacterial (n = 17) genomes, archaeal (n = 3), fungal (n = 3), and viral (n = 3) species (total species = 26) were included. Genomes were selected for strains which originate from the mammalian gut. Only the nuclear genomes of the fungi were included.

The under- and overprediction of the tools was quantified using ORForise^52^, which provides metrics on the performance of gene prediction software, by comparing predicted results to reference annotations (**Supplementary Table 3**).

### Datasets

A total of 9,677 metagenomes from 43 studies were analysed (**Supplementary Table 4**). Among those, 7,735 metagenomic assemblies from 41 studies were available from Pasolli *et al.* (2019)^8^ and 1,326 assemblies from the Integrative Human Microbiome Project^9^. FASTQ files for 616 metagenomes from a Japanese cohort^53^ were also downloaded and processed. Human sequences were removed from the latter metagenomic reads by running BBMap^54^ (v38.18) with Genome Reference Consortium Human Build 38 (GRCh38)^55^ as a reference. Reads were assembled using MEGAHIT^56^ (v1.2.9) with a range of k-mer sizes from 21 to 99 and a minimum count of 5 to filter out low-quality reads. Metagenomes with less than 1,000 proteins predicted were removed (n = 43), meaning 9,634 metagenomic samples were retained. Metadata for metagenomic samples was either obtained directly, or the curatedMetagenomicData R package^57^. Additionally, a collection of 3,594 non redundant high-quality genomes from Leviatan *et al.* (2022) were studied, this included both metagenomic assembled genomes (MAGs) and isolate genomes. These non-redundant genomes had a reported median completeness of 95% and median contamination of 0.67%^58^.

### Taxonomic assignment

For the taxonomic classification of all contigs, Kraken 2 (v2.1.2) was applied with a confidence threshold of 0.15^59^. To enhance the taxonomy-assigned fraction of each metagenome, a custom Kraken 2 database was formed using all complete genomes for Archaea, Bacteria, Eukaryota, and Viruses from the NCBI RefSeq database (April 2023)^60^. The inclusion of the human genome (Genome Reference Consortium Human Build 38 (GRCh38)) within the Kraken 2 database facilitated the removal of host contamination from downstream gene prediction. Based on the taxonomic ID assigned to each contig, the correct genetic code was assigned according to the NCBI Taxonomy Database (August 2023)^61^. For representative genomes, the taxonomic classifications of Leviatan *et al.* were computed with GTDB-Tk (v2.3.2; r214)^58,62^. Contigs from each metagenome/genome were subdivided into files based on their inferred taxonomy and genetic code with the selected combination of three gene prediction tools being run on the corresponding subset files. Contigs without an assigned taxonomy were annotated with the bacteria-optimised selection of tools and genetic code 11.

### Gene prediction

Variability in tool outputs was accounted for with a script to parse the start, stop, and strand information of prediction outputs. When multiple tools predicted genes with the same stop position, but different start positions, the longest sequence was selected. In cases of differing stop codons, all sequences were retained. Secondly, all partial genes were removed from consideration, including those with a start codon in the first or last three bases of a contig, where assembly errors are common^63^, to prevent the inclusion of truncated genes. When possible, the minimum prediction length setting was set to 21 bases (six amino acids plus stop codon).

Based on this information, protein sequences were generated using the genetic code corresponding to the Kraken 2 taxonomy ID predictions. No protein count threshold was applied for the number of predicted genes in the studied representative genomes.

### Protein ecology across human gut samples (InvestiGUT)

InvestiGUT (v0.1) accepts either a single protein sequence or a set of sequences, as defined by the options ‘–s’ or ‘–m’ respectively. User-provided protein sequences are matched to the collection of human gut microbiome proteins via DIAMOND (v2.1.8.162)^64^ alignment with a default minimum query and subject coverage of 90% and identity of 90%, which can be defined by the user. User proteins can be analysed individually (-s), or as a group (-m) where all proteins must be present within a sample to be considered. The group analysis allows users to determine how frequently all enzymes in a certain pathway or all subunits in a protein complex are present. Output is divided into metagenomic and species-based analysis results, with both being available as raw data in the form of TSV files and as automatically generated vector figures.

Metadata for each metagenomic sample included age, gender, BMI, smoker-status, westernised-diet status, use of antibiotics, and geographical location. Additionally, disease prevalence includes output for Crohn’s disease, ulcerative colitis, colorectal cancer, *Clostridioides difficile* infection, type-2 diabetes, rheumatoid arthritis, and fatty liver disease. Metadata vocabulary is consistent with that used in the curatedMetagenomeData repository. Prevalence of the queried sequence in each of these groups is quantified, and compared either between categories, or healthy controls, using Fisher’s exact test from the scipy.stats^65^ module in Python.

Querying the user input protein against the 3,594 nonredundant genomes provides a list of species which contain the protein, termed “function-positive species”. The taxonomic range of the protein is determined by the prevalence of the protein with each genus, family, and phylum of positive species. The function-positive fraction is obtained as the cumulative relative abundance of function-positive genomes across 4,624 individuals for which pre-computed relative abundance values are available. InvestiGUT is available at: https://github.com/Matt-Schmitz/InvestiGut

### Gene catalogue creation

The protein sequences from the 9,634 filtered human gut metagenomes and 3,594 genomes were clustered using MMseqs2 (v14-7e284) linclust using options ‘--cov-mode 1 -c 0.8’ (minimum coverage threshold of 80% the length of the shortest sequence), ‘--kmer-per-seq 80’ (number of k-mers selected per sequence), and ‘--min-seq-id 0.9’ to cluster at 90% protein identity. Thresholds were selected as previously used to generate the UHGP-90^66^. Overlap between the MiProGut-90 and UHGP-90 was identified by combining the proteins of both catalogues and reclustering with the same options.

### Functional annotation

The MiProGut gene-catalogue was annotated using eggNOG-mapper (v2.1.12) with the inbuilt DIAMOND blastp search option and default parameters^67,68^. Further annotation was conducted with MANTIS using the integrated annotation file^69^. When proteins had more than one functional category or general category letter association, the corresponding functional descriptions of each category were evenly weighted. Antimicrobial activity was predicted using the DBAASP toolkit^70^. Positive activity against *Escherichia coli* ATCC 25922, *Pseudomonas aeruginosa* ATCC 27853, *Klebsiella pneumoniae*, *Staphylococcus aureus* ATCC 25923, *Candida albicans*, and *Saccharomyces cerevisiae* was determined as a predicted MIC <25 μg/ml. Annotation against the NCBI nr database was conducted using BLASTP with default options^71^.

### Metatranscriptome expression analysis

DIAMOND (v2.1.8.162)^64^ mapped metatranscriptomic reads^7,72,73^ against the MiProGut-90 and UHGP-90 protein catalogues using the ‘blastx’ command with options ‘--id 90 --evalue 1e-6 -k 1 --max-hsps 1’. The aligned fraction was calculated per-sample as: (aligned reads / total reads) × 100. Expression of specific proteins was quantified as reads per kilobase per million mapped reads (RPKM) and calculated using: (aligned reads / ((gene length / 1,000)) x (total reads / 1,000,000).

## Results

### Lineage-specific gene prediction expands the human gut protein repertoire

A workflow for lineage-optimised gene prediction was developed using targeted selection of gene prediction tools based on the taxonomic assignment of each metagenomic sequence and customisation of each tool’s options (genetic code, gene size) (**Figure 1a**). The selection of tools for gene prediction was informed by initial testing of 13 tools on archaeal (n = 3), bacterial (n = 17), fungal (n = 3), and viral (n = 3) species (total species = 26). The quality of each tool’s annotations was quantified and compared using ORForise^52^. Given the variability of individual tools to predict all genes, we investigated the potential synergy of combining two or three tools in tandem to predict genes (**Supplementary Table 3**). Given the small, but consistent increase in both full and partial genes predicted by the combination of three tools, and the low cost in terms of spurious genes compared to two tools, we chose to use the combination of three tools that performed best for each taxonomic group.

**Figure 1:**
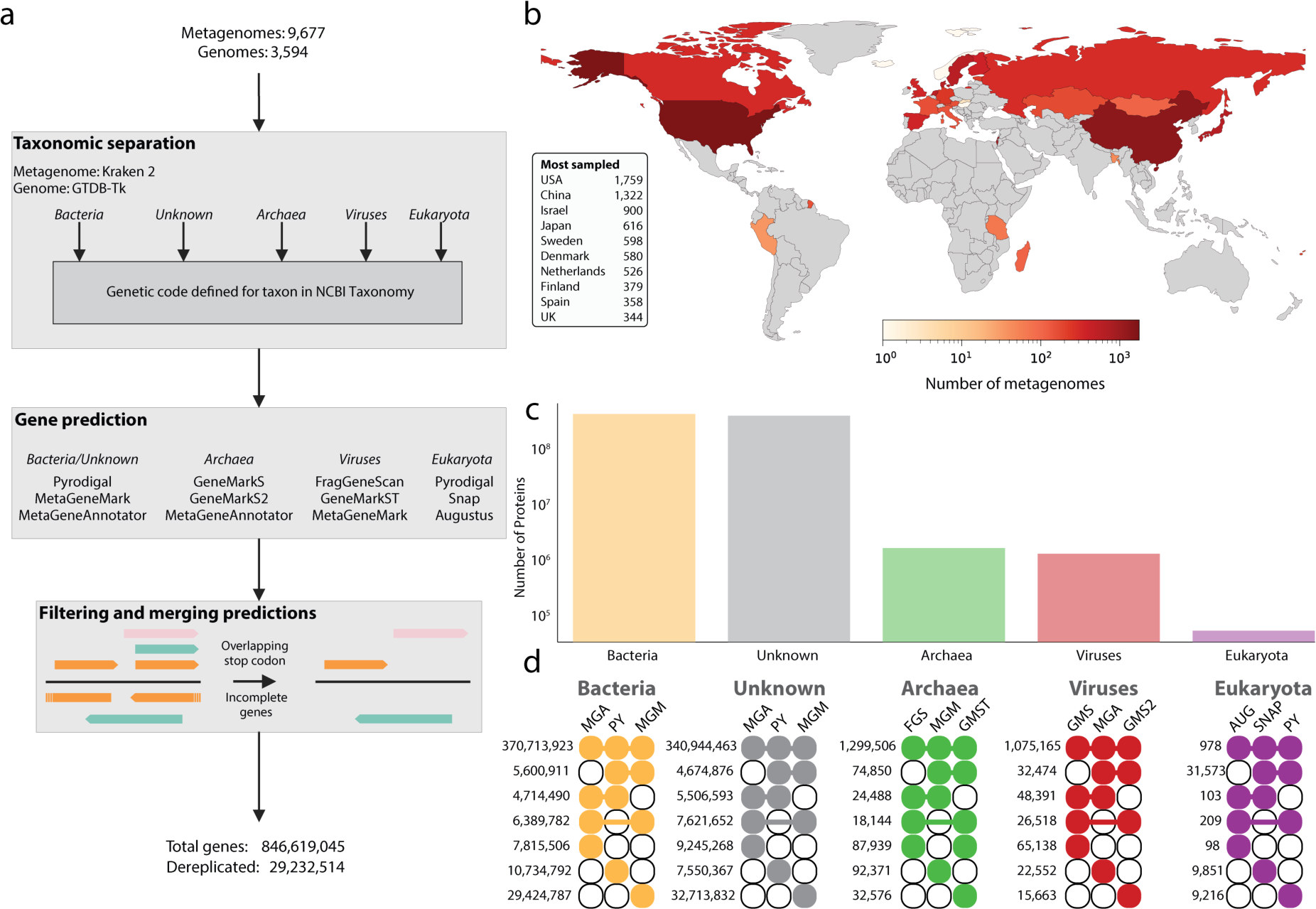
Taxonomy-informed gene prediction workflow and its application to the human gut. **a:** The workflow consists of three major steps; Taxonomic separation of either metagenomic contigs, or input genomes, into their respective domains. The genetic code utilised by each lineage is then identified at the family level, to account for variation within domains of life. For each domain-genetic code grouping, the respective gene prediction tools are run, and then merged, removing redundant predictions and incomplete gene predictions occurring at the edges of contigs. **b**: Geographical distribution of the human gut metagenomes used, with darker colours indicating a greater number of samples originate from the country. **c**: The number of proteins predicted for each taxonomic domain **d**: The overlap between gene prediction tools was identified for each domain to quantify each tool’s variation from consensus predictions. Visualisation is based on a variation of a upset plot, where the number on the right is the number of overlapping protein predictions from the tools indicated by the connected coloured circles. Abbreviations for tools are: MGA = MetaGeneAnnotator; MGM = MetaGeneMark; FGS = FragGeneScan; GMS = GeneMarkS; GMS2 = GeneMarkS2; GMST = GeneMarkST; PY = Pyrodigal; SNAP = SNAP, AUG = AUGUSTUS.

This workflow was applied to 9,677 metagenomes from 28 countries (**Figure 1b**). In addition to the metagenomes, a non-redundant collection of genomes representing the prokaryotic diversity within the human gut was included for downstream analysis of the taxonomic occurrence of proteins^58^. Taxonomic profiling of the metagenomic samples was consistent with previous observations that identified Bacteria as the dominant microbial domain in the human gut (**Supplementary Table 5**). The predicted proteins were dominated by those originating from bacterial contigs (58.4 ± 18.9%), followed by proteins on contigs that could not be assigned to a specific domain by Kraken 2, termed unknown (41.2 ± 18.8%), then viruses (0.19 ± 0.409%), archaea (0.154 ± 0.645%), and eukaryotes (0.0320 ± 1.31%) (**Supplementary Table 6**). The high percentage of taxonomically unassigned proteins, the unknown group, is in line with current estimates that >50% of the gut microbiota has yet to be cultured, hence complete genomes, which were used for taxonomic assignment, do not exist for many gut microbes^74,75^.

In total, 846,619,045 genes (metagenomes: 838,528,977, genomes: 8,090,068) were predicted, with the majority originating from contigs of bacteria or unknown assignment (**Figure 1c**). In comparison, the exclusive use of Pyrodigal across all metagenomes identified 737,874,876 genes (metagenomes: 730,237,038, genomes: 7,637,838), meaning that the lineage-specific workflow identified an additional 108,744,169 genes (14.7%). While the use of multiple gene prediction tools might be expected to inflate the number of predicted genes, the majority of predictions were consistent across tools, except for eukaryotes. For eukaryotic contigs, AUGUSTUS^30^ resulted in substantially lower gene prediction numbers, reducing the harmonious prediction of eukaryotic genes across all three tools (**Figure 1d**). In addition to the lower number of genes predicted by AUGUSTUS, the overlap between SNAP and Pyrodigal also highlights inconsistency. This is likely due to the inability of Pyrodigal to predict multi-exon genes, whereas both SNAP and AUGUSTUS predict exons and introns, which is crucial for accurate gene prediction in eukaryotes^76^.

### Enhanced coverage of the protein landscape from the human gut

Lineage-specific gene prediction yielded a larger number of proteins compared to any single approach. Therefore, we sought to confirm that these were real proteins and not spurious predictions. To facilitate comparison with a previously established protein catalogue, the Unified Human Gastrointestinal Protein (UHGP) catalogue^66^, we dereplicated our >800 million proteins at 90% similarity to 29,232,514 protein clusters, increasing the human gut protein landscape by 210.2% compared to UHGP. We termed this protein catalogue the Microbial Protein Catalogue of the Human Gut (MiProGut).

Nearly 10,000 samples were used to generate MiProGut, but the rarefaction of MiProGut suggests that greater diversity has yet to be captured for all taxonomic groups (**Figure 2a**). This was supported by analysis of 100 permeations where 2,397 samples were needed to cover 50% of proteins in MiProGut, and for 90% coverage an average of 7824 samples were needed. Eukaryotic proteins were observed least frequently, with the majority of these proteins being identified within a few samples that were dominated by Eukaryotic contigs (**Supplementary Table 5**). The Western bias of included samples, as well as low number of samples from developing countries may explain the rarefaction observation of further diversity. The inclusion of additional samples may also reduce the number of protein clusters consisting of a single protein sequence, known as singletons (n = 14,043,436). This frequency of singleton clusters is consistent with previous observations that most protein clusters are rare^77^, leading to the formation of singletons. While singletons were rarely captured by metagenomic sequencing, expression in human gut samples was observed for 39.1% of singletons, confirming they are not spurious and are functionally relevant to the microbiota.

**Figure 2:**
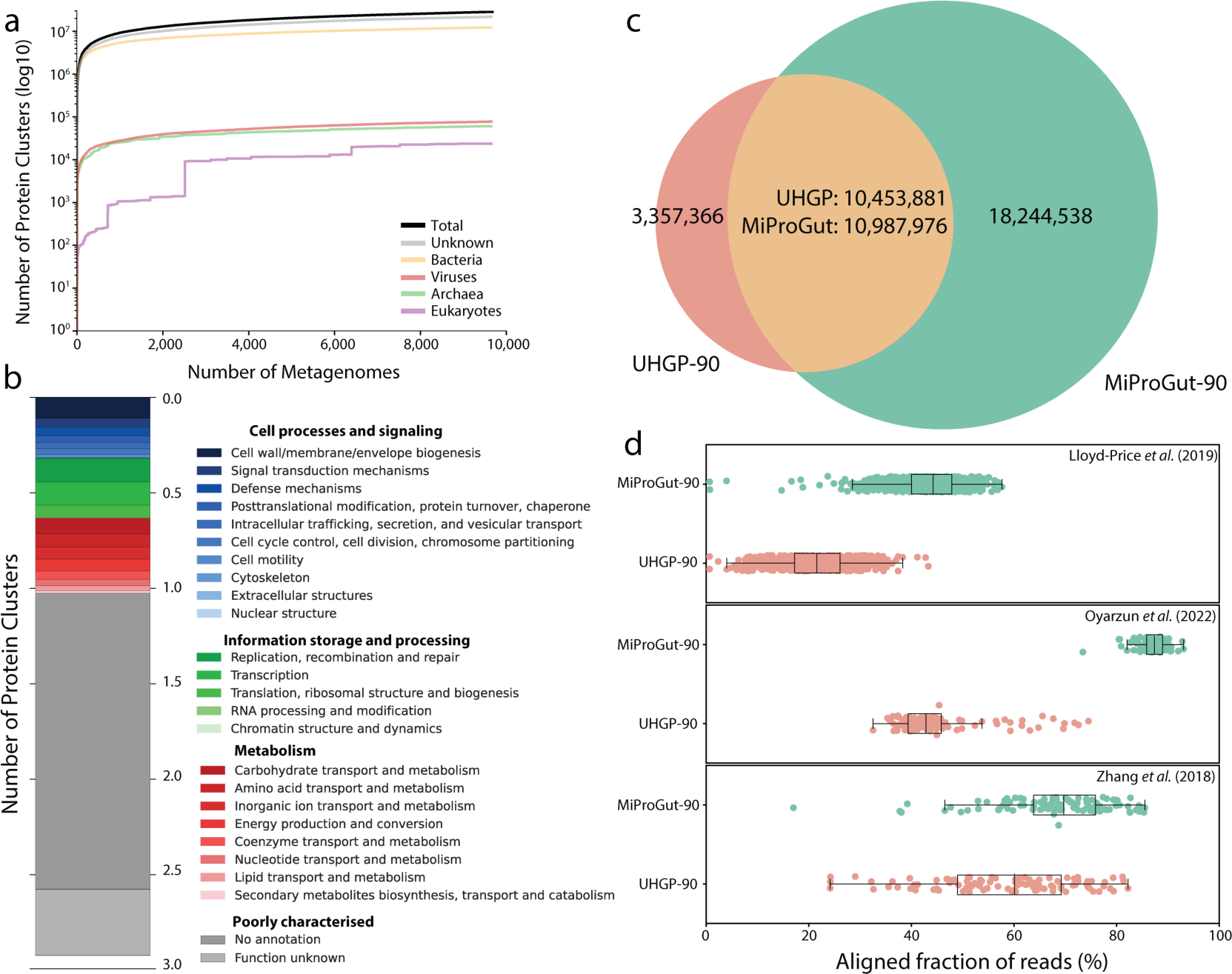
Taxonomy-informed gene prediction expands the protein landscape of the human gut microbiome. **a**: For each taxonomic group, as well as the cumulative protein repertoire of the of proteins within the MiProGut (log10 scale), refraction analysis was conducted across all studied metagenomes. **b**: Functional annotation of the MiProGut based on COG functional groupings. **c**: Comparison of MiProGut (green) to UHGP (orange), both clustered at 90% amino acid identity. **d**: The aligned fraction of metatranscriptomic reads from multiple human gut studies (Lloyd-Price *et al.* 2019: n = 682, Oyarzun *et al.* 2022: n = 100, Zhang *et al.* 2018: n = 100) by both MiProGut and UHGP.

MiProGut provides an improved resource for protein sequence identification, but functional annotation is required to understand the role of these proteins, both in relation to the microbiome, and their impact on the host^78,79^. The majority (64.4%) of MiProGut proteins lacked informative functional annotation when annotated with eggNOG-mapper (**Figure 2b**). Further annotation with MANTIS, which integrates multiple functional databases, increased the annotated fraction by 4.9% compared to all eggNOG assigned COG categories. However, this included assignments to general functions. These values re-highlight the high number of microbial proteins in the human gut that lack meaningful functional annotation^74^.

Significant overlap was observed between MiProGut and UHGP, with 75.7% of UHGP proteins being covered by MiProGut, whereas only 37.6% of MiProGut proteins were covered by UHGP (**Figure 2c**). Compared to UHGP, MiProGut was enriched for eukaryotic functions, especially nuclear structure proteins (3.5 Vs 227.3 protein clusters respectively).

Identification of overlapping proteins between the two catalogues provides support that these proteins are not spurious. Expression was quantified in 862 metatranscriptomic samples from three studies (**Figure 2c**)^7,72,73^. We identified transcriptional evidence for 13,756,670 MiProGut proteins (47.1%), compared to 7,689,106 UHGP proteins. Given the increased number of expressed proteins, we calculated the aligned fraction of reads for each of the 862 metatranscriptomes. Across all samples, MiProGut covered an additional 22.9 ± 9.9% reads compared to UHGP. The largest increase occurred in a Spanish cohort where MiProGut increased read coverage by an additional 41.6 ± 8.6%^72^.

### Identification of commonly expressed small protein clusters within the human gut

Small proteins have been understudied in the human gut as many gene prediction tools exclude them by default^80^. To improve the prediction of these proteins, we modified the parameters to predict proteins of >5 amino acids. This resulted in the prediction of 44,164,853 small proteins (5.2% of total proteins), represented by 3,571,095 protein clusters within MiProGut, called small protein clusters (SPCs) (**Figure 3a**). Of these, 1,104,393 (30.9%) clusters were singletons (**Figure 3b**), while the largest SPC contained 10,700 proteins and was identified as a member of the proposed ‘family 350024’ of crosstalk proteins^80^. Studying the expression of the SPCs within 687 metatranscriptomic samples revealed that, while many were rarely expressed (<10% of samples), 69 SPCs were highly prevalent, being expressed in >90% of samples (**Figure 3c**).

**Figure 3:**
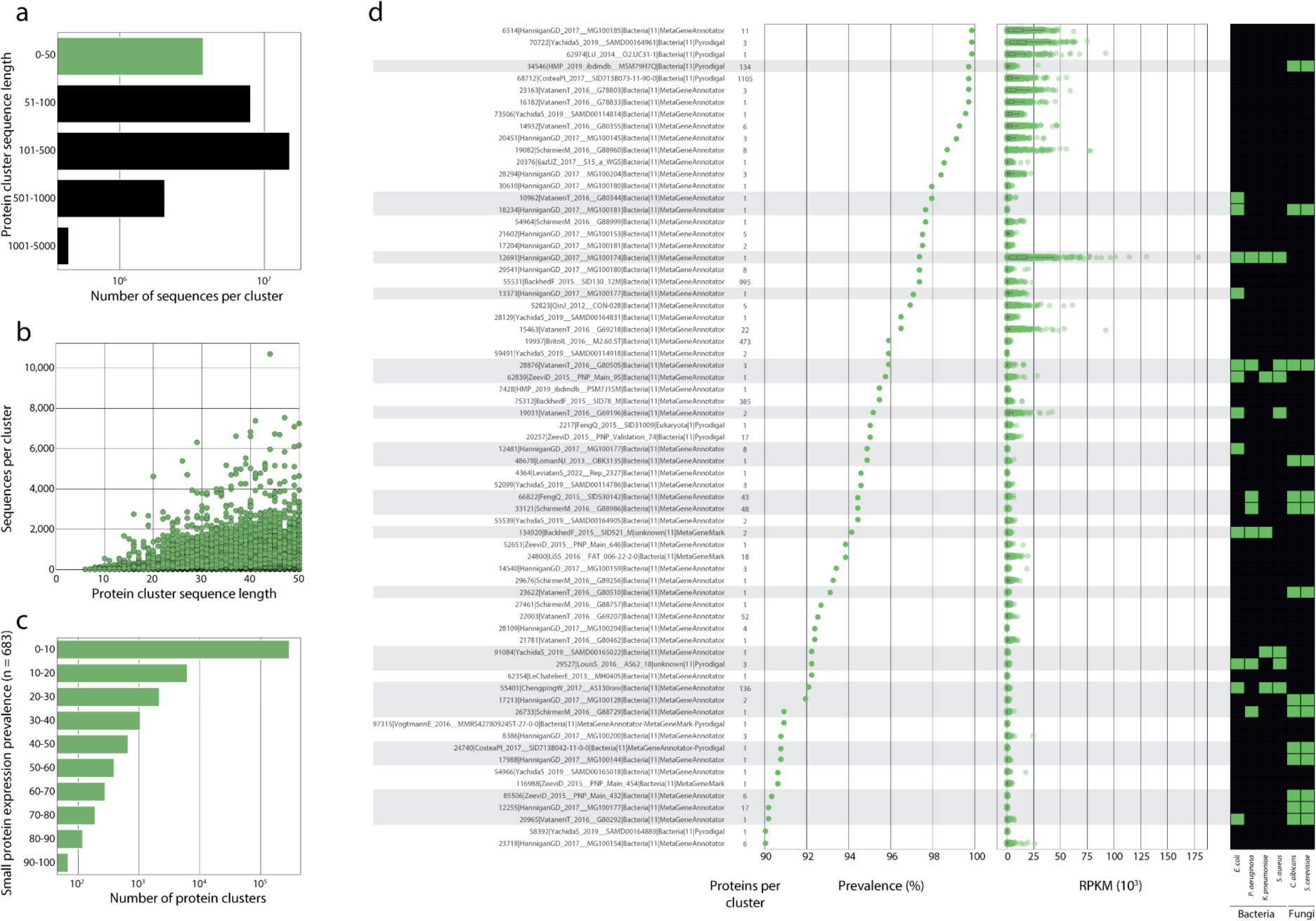
Identification of highly prevalent, and expressed small proteins within the gut and their antimicrobial potential. **a**: The size of protein cluster representative sequences, with small proteins (≤50 amino acids) in green. **b**: The number of members within each protein cluster, plotted against the size of each protein clusters representative sequence. **c**: The prevalence of each protein clusters expression across the 687 metatranscriptomic samples from Lloyd-Price *et al.* (2019)^7^. **d**: The name, number of member sequences, expression prevalence, RPKM and predicted antimicrobial activity for proteins expressed within ≥90% of metatranscriptomic samples (n = 69).

The gastrointestinal tract has been identified as a rich source of small proteins with antimicrobial activity^81,82^. To determine whether this included the most commonly expressed SPCs, we predicted each SPCs antimicrobial activities^83^ (**Figure 3d**). By predicting the antimicrobial activity against four ESKAPE pathogens and two fungi of relevance to human health, we found that 35% (24/69) of these SPCs could be antimicrobial. The most highly expressed antimicrobial SPC (13,707 ± 17,886 RPKM) was predicted to be active against all four ESKAPE pathogens (*Escherichia coli* ATCC 25922, *Pseudomonas aeruginosa* ATCC 27853, *Klebsiella pneumoniae*, *Staphylococcus aureus* ATCC 25923). Although expressed in 97.1% of the samples, this cluster contained only a single sequence with similarity to an unknown protein from *Bifidobacterium* spp. The coding of this protein by a dominant genus of human gut commensals, in addition to its high expression and wide range of predicted activity, suggests this protein may be of importance to the gut ecosystem. Of the 24 antimicrobial SPCs, 14 were predicted to target both fungal species, *Candida albicans* and *Saccharomyces cerevisiae*.

### Ecological distribution of proteins across individuals uncovers associations with host health status

The application of lineage-specific gene prediction enhances the ability to study the ecological distribution of proteins, referred to in this manuscript as protein ecology. Using the metadata associated with both the metagenomes studied and the genomes analysed, we created a tool, InvestiGUT, to facilitate protein ecology studies of the human gut (**Figure 4a**).

**Figure 4:**
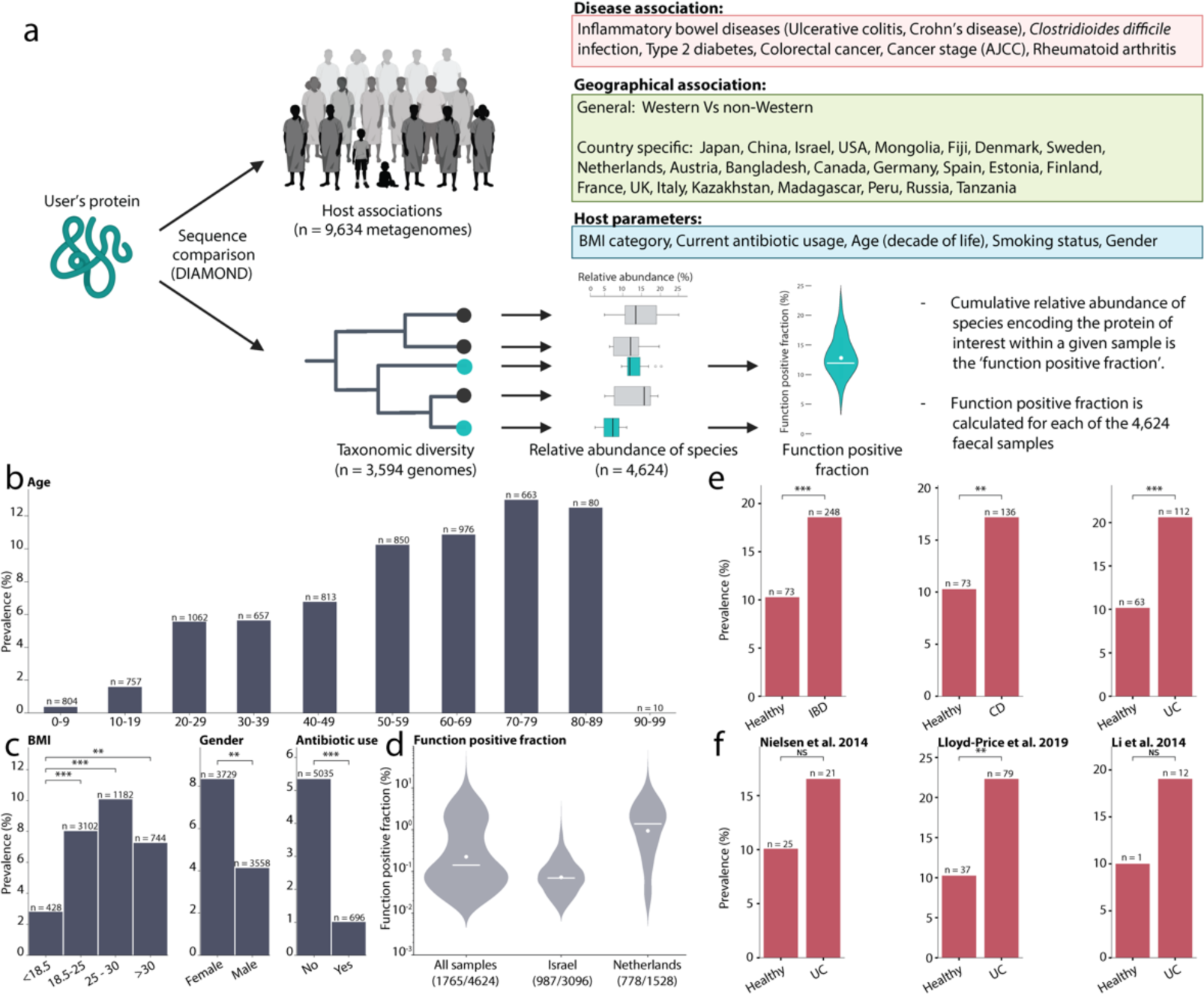
Application of protein ecology to the human gut identifies associations between microbial functions and host parameters. **a**: Workflow of InvestiGUT, detailing the host associations that are studied, as well as detailing the process for calculation of the function positive fraction. Functional positive species and their relative abundances are coloured turquoise, along with the final functional positive fraction. Methyl-co-reductase was used as a use-case of the multiple-protein option within InvestiGUT, identifying the complexes prevalence across host age (**b**), BMI, gender, and antibiotic use (**c**), and determine the MCR function positive fraction of the microbiome from 4,624 individuals (**d**). As a second use-case, the small antimicrobial peptide ‘55401|ChengpingW_2017 AS130raw|Bacteria|11|MetaGeneAnnotator’ was studied. Significant associations with IBD, CD, and UC were observed across studies (**e**). To explore the association with UC further, the three individual studies into UC were examined, each showing increased prevalence in UC patients compared to healthy controls (**f**). Significance from Fisher’s exact test, after Benjamini-Hochberg correction, are shown as: NS = not significant, * < 0.05, ** < 0.01, *** < 0.001.

InvestiGUT was developed to accept multiple sequences at once and examines them either individually or as a collection, where only samples that are positive for all queried sequences are reported. The input sequences are compared against all 846,619,045 predicted protein sequences, representing those from both the metagenomes and genomes. The sequence similarity for annotation is by default 90%, but can be defined by the user. Integration of metadata for the metagenomes is used to quantify the prevalence of the queried proteins with disease and geographic locations, as well as host parameters. The inclusion of genomes from human gut commensals allows the taxonomic range of the protein to be determined. Additionally, we can determine the ‘function positive fraction’, which represents the proportion of an individual’s microbiota that is positive for the function/protein sequence of interest. This is calculated by determining which genomes encode the protein sequence of interest and then determining the cumulative relative abundance of these genomes across 4,464 individuals^58^.

Methanogenesis in the human gut is restricted to archaea and has been linked to several human diseases^84^. Methyl-coenzyme M reductase (MCR) catalyses the formation of methane in methanogens and consists of an alpha, beta, and gamma chain. Due to MCRs multiple chains, the multiple sequence approach was used in InvestiGUT, ensuring only those samples in which all three chains were detected to be studied. The prevalence of methanogenic functionality increased with host age (**Figure 4b**), confirming previous reports of this association^85^. Increased methane production has been reported in individuals with higher BMI, supporting our observation that MCR was more prevalent in individuals with higher BMI^86^ (**Figure 4c**). Variation with gender has also been previously observed, although age is a confounding variable in this association^85^. The decreased prevalence of MCR in patients noted as having used antibiotics provides the first metagenomic insight into the observation that antibiotic therapy can eliminate methane in the breath of individuals^87^. This may be due to indirect interactions, such as the dependence of methanogenic archaea on bacteria that are themselves susceptible to the antibiotics. The MCR positive fraction revealed a country-specific association, with individuals from the Netherlands having a higher relative abundance of species positive for MCR than individuals from Israel (**Figure 4d**).

Country-specific associations can be explained by variation in diet. For example, the enrichment prevalence of seaweed-degrading porphyranases (*Bp1689*), and agarases (*Bp1670*) in the gut of Japanese individuals, compared to North Americans^2^. The original study was limited to 31 individuals (13 Japanese vs 18 North Americans), hence our analysis expands the study of these proteins across 28 countries. We confirmed that both proteins are most prevalent in Japanese samples (56.8% and 79.4% respectively), followed by China (18.5% and 27.4% respectively) (**Supplementary Figure 1**). Interestingly, 41 USA samples were positive for *Bp1670* and 39 for *Bp1689*, perhaps reflecting changes in diet, travel, or immigration over the last decade.

Given that small proteins are highly expressed in the human gut, we investigated the ecological distribution of the 69 most highly expressed SPCs. Of these, we observed that 7 were enriched in inflammatory bowel disease (IBD), 9 in ulcerative colitis (UC), and 8 in Crohn’s disease (CD). In particular, ‘55401|ChengpingW_2017 AS130raw|Bacteria|11|MetaGeneAnnotator’ showed an significant change in prevalence with all three conditions across the studies (**Figure 4e**). When we explored this association further, we found that the association with UC was only significant in one of the three studies, although the increased prevalence in UC patients was observed in all three studies (**Figure 4f**). Analysis of the 69 SPCs expression in patients with UC and Crohn’s disease CD patients compared to control patients identified that 31 and 34 proteins respectively were enriched in IBD sub-types, including ‘55401|ChengpingW_2017 AS130raw|Bacteria|11|MetaGeneAnnotator’ (**Supplementary Table 7, 8**). This association may be explained by the predicted antimicrobial activity of the protein, which targets *E*. *coli*^88^, *K*. *pneumoniae*^89^, and *S*. *aureus*^90^, all of which have been studied as causative agents of colitis.

The largest SPC (n = 10,700) was also of interest due to its prevalence. It contained a highly conserved core sequence across all sequences, forming a continuous helix (**Supplementary Figure 2a**). The taxonomic range of this small protein was restricted to the *Bacteroidaceae*, in which it is prolific, with 76.3% (45/59) of *Bacteroides* spp. containing a matching protein sequence (**Supplementary Figure 2b**). Functional annotation of the protein suggested that it has antimicrobial activity against *S. aureus* (**Supplementary Figure 2c**). The prevalence of this protein was associated with Westernised individuals and decreased with antibiotic use (**Supplementary Figure 2d**). These use cases highlight InvestiGUT’s ability to facilitate protein ecology studies, confirm associations previously identified in smaller cohorts, and identify associations uncovered by integrating multiple data sources.

## Discussion

Integrating the taxonomic assignment of a sequence into the choice of gene prediction tools, and optimising the parameters used, revealed millions of protein sequences previously missing from human gut gene catalogues. These included proteins specific to groups that have previously been overlooked, such as the eukaryotes, archaea and viruses^28^. The lack of functional annotation by both MANTIS and eggnog-mapper for 47.9% of the proteins is consistent with previous observations that nearly half of the identified proteins in the human gut have yet to be characterised^74^. Integration of metatranscriptomic datasets facilitated the identification of highly expressed small proteins with potential antimicrobial activity against key human pathogens. Further analysis of these proteins revealed that the majority (49/69) were differentially expressed between IBD subtypes and non-IBD patients, including those with predicted antimicrobial activity. The association with human health conditions, as well as their antimicrobial activity, warrants further investigation of these proteins to characterise their impact on the microbiota and potential therapeutic application.

The increased coverage of the human gut protein landscape by MiProGut, compared to other gene catalogues, facilitates greater analysis of omics datasets, doubling the aligned fraction in a metatranscriptomic study. As gene catalogues are often used as reference databases for the metagenomic study of an environment, the discovery of these missing proteins will facilitate the identification of disease-specific biomarkers that were previously overlooked. However, the bias of metagenomic samples included towards Westernised countries (USA, China) suggests that the human gut landscape could be further enhanced with the inclusion of additional samples from underrepresented countries and age groups (infants, elderly)^21^.

We present the concept of protein ecology, which focuses on the study of proteins rather than taxonomic species and should be applied more broadly to the study of the human gut. InvestiGUT facilitates protein ecology studies by providing statistical analysis of the prevalence of a protein sequence/sequences of interest across the samples examined in this work. The application of this approach identified both previously observed associations, but across a larger number of samples, providing independent validation of these associations. In addition to protein prevalence, the relative abundance of microbial species containing the queried protein, referred to as the ‘function positive fraction’, is of interest for protein ecology, as it allows the estimation of protein distribution within the ecosystem. The current genome collection and approach does not account for strain-level diversity, instead assuming that the presence within the reference genome is representative. Currently, this limits our ability to truly capture the gut microbiotas diversity. Future integration of a larger collection of genomes that captures the strain-level diversity of each species would facilitate the calculation of a modified functional positive fraction.

## Data availability

Gene and protein predictions, along with the nonredundant MiProGut catalogue, are available at: https://zenodo.org/doi/10.5281/zenodo.10988030. InvestiGUT is available at: https://github.com/Matt-Schmitz/InvestiGut

## Funding

TCAH received funding from the RWTH Start-up grant, titled ProtoBIOME and University Hospital of RWTH Aachen START grant, titled LeakyGut (021/23). TC received funding from the German Research Foundation (DFG): project no. 403224013 – SFB1382 Gut-liver axis, subproject Q02; project no. 460129525 – NFDI4Microbiota.

## Acknowledgements

The authors would like to thank Susan Jennings and Charlie Pauvert for reviewing the paper prior to submission, and Amy Coates for designing the logo.

## Supplementary Tables

**Supplementary Table 1: Gene Prediction Software.** For each gene prediction tool studied, the version, target taxa, and reference are provided.

**Supplementary Table 2: Species studied for optimisation of gene prediction.** For each species studied, the genome name, domain, and database are defined. Those species placed in the ‘probiotic’ and ‘pathogen’ groups are also defined.

**Supplementary Table 3: Comparison of gene prediction tools.** For each genome assigned to the studied domain (Archaeal, Bacterial, Eukaryotic, Viral) tools were tested individually, in pairs, and triplets. The perfect, partial, and spurious gene predictions as determined by ORForise are reported.

**Supplementary Table 4: Human gut metagenomic studies analysed.** For metagenomic study used to create MiProGut and the InvestiGUT database, the ‘curated metagenomic data’ identifier, number of included metagenomes (post filtering), and the studies PubMed identifier are provided.

**Supplementary Table 5: Taxonomic assignment of metagenomic contigs.** The number of contigs assigned to Archaea, Bacteria, Eukaryota, Viruses, or defined as unknown are provided, both for each individual sample, and grouped by study.

**Supplementary Table 6: Taxonomic assignment of predicted proteins.** The number of proteins predicted from contigs assigned to each taxonomic group is stated for each metagenomic study.

**Supplementary Table 7: Differential expression analysis of frequently expressed small proteins between ulcerative colitis and healthy samples.** For each of the 69 small protein clusters expressed within at least 90% of studied metatranscriptomic samples, the RPKM within ulcerative colitis and healthy samples was calculated and compared. Significantly differentially expressed (adj. P-value < 0.05; Benjamini-Hochberg correction) protein clusters are reported along with their mean RPKM within each group and log2 fold change between groups.

**Supplementary Table 8: Differential expression analysis of frequently expressed small proteins between Crohn’s disease and healthy samples.** For each of the 69 small protein clusters expressed within at least 90% of studied metatranscriptomic samples, the RPKM within Crohn’s disease and healthy samples was calculated and compared. Significantly differentially expressed (adj. P-value < 0.05; Benjamini-Hochberg correction) protein clusters are reported along with their mean RPKM within each group and log2 fold change between groups.

## Supplementary Figures

**Supplementary Figure 1:**
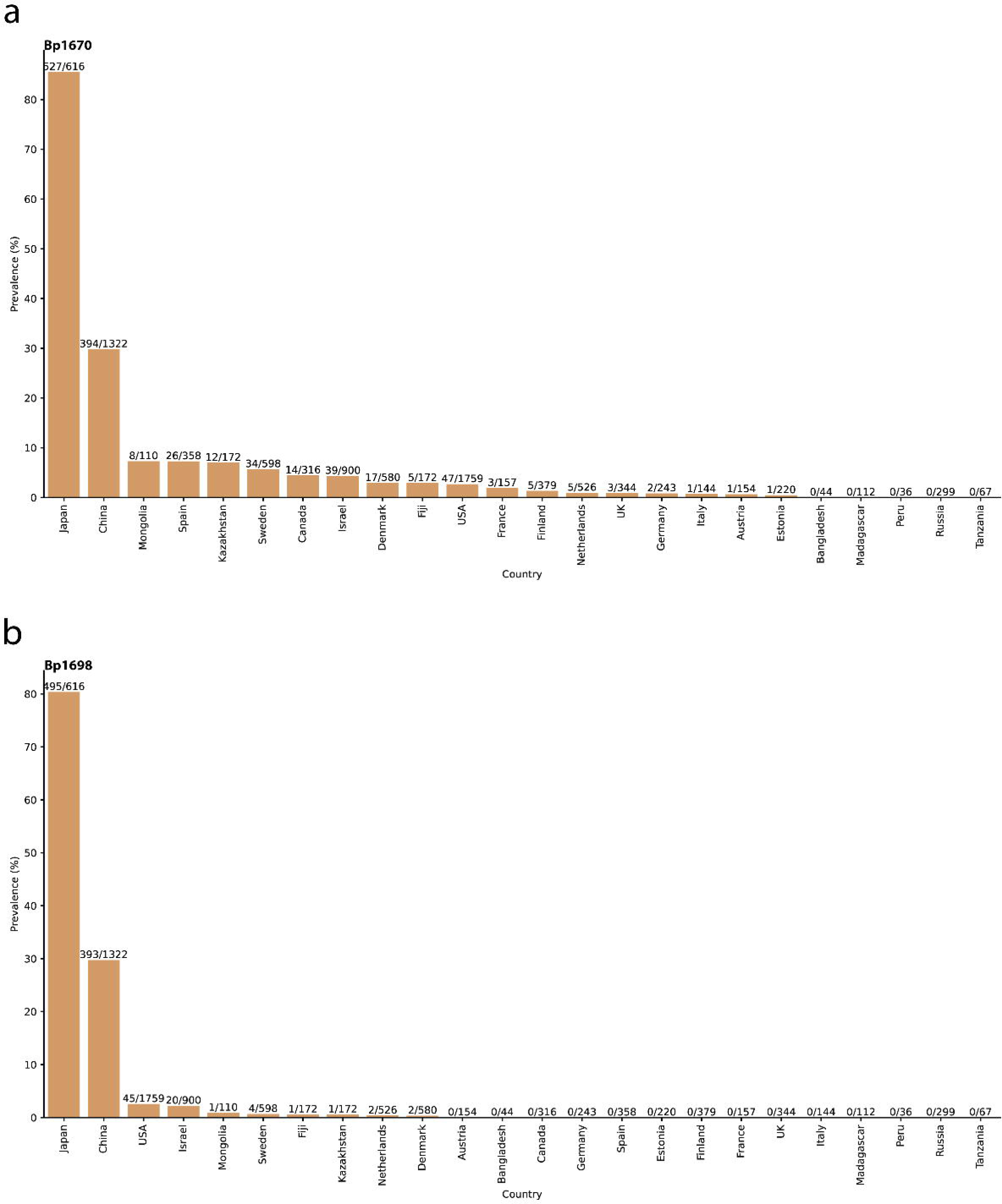
Geographical prevalence of seaweed degrading porphyranases (*Bp1689*), and agarases (*Bp1670*) Each protein was studied using InvestiGUT at the default 90% identity threshold.

**Supplementary Figure 2:**
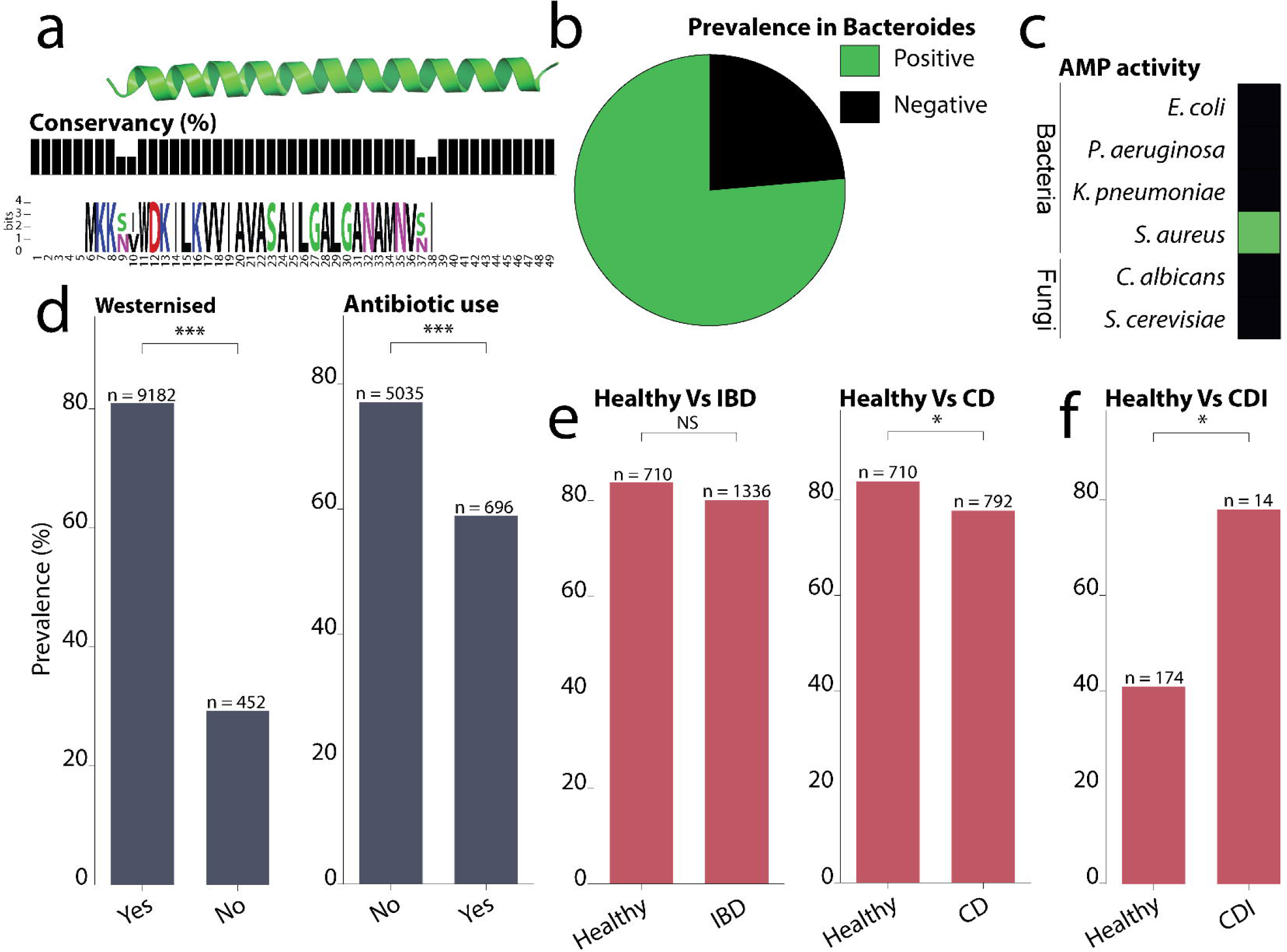
Analysis of the small protein cluster ‘33977|YachidaS_2019 SAMD00114748|Bacteria|11|MetaGeneAnnotator-MetaGeneMark-Pyrodigal’. The conservation of sequence between members of the cluster, along with an Alphafold2 structural model were generated (**a**). The frequency of the cluster across *Bacteroides* spp. (n = 59) (**b**), and predicted antimicrobial activity against key pathogens and fungi (**c**). The prevalence of the cluster with host parameters, including westernised status, and antibiotic use, was studied (**d**), along with the prevalence in IBD and Crohn’s disease (CD) patients (**e**), and *Clostridioides difficile* infection (CDI) patients compared to healthy controls (**f**). Significance from Fisher’s exact test, after Benjamini-Hochberg correction, are shown as: NS = not significant, * < 0.05, ** < 0.01, *** < 0.001.

## Notes

### Competing Interest Statement

The authors have declared no competing interest.

